# Clarifying Microbial Nitrous Oxide Reduction Under Aerobic Conditions: Tolerant, Intolerant, and Sensitive

**DOI:** 10.1101/2022.10.27.514152

**Authors:** Zhiyue Wang, Nisha Vishwanathan, Sophie Kowaliczko, Satoshi Ishii

**Affiliations:** BioTechnology Institute, University of Minnesota, St. Paul, MN, USA; Department of Soil, Water, and Climate, University of Minnesota, St. Paul, MN, USA

**Keywords:** Nitrous oxide reduction, oxygen sensitivity, microsensor, biokinetics

## Abstract

One of the major challenges for the bioremediation application of microbial N_2_O reduction is its oxygen sensitivity. While a few strains were reported capable of reducing N_2_O under aerobic conditions, the N_2_O reduction kinetics of phylogenetically diverse N_2_O reducers are not well understood. Here we analyzed and compared the kinetics of Clade I and Clade II N_2_O-reducing bacteria in the presence or absence of O_2_ by using a whole-cell assay with N_2_O and O_2_ microsensors. Among the seven strains tested, N_2_O reduction of *Stutzerimonas stutzeri* TR2 and ZoBell were not inhibited by oxygen (i.e., oxygen tolerant). *Paracoccus denitrificans, Azospirillum brasilense*, and *Gemmatimonas aurantiaca* reduced N_2_O in the presence of O_2_ but slower than in the absence of O_2_ (i.e., oxygen sensitive). N_2_O reduction of *Pseudomonas aeruginosa* and *Dechloromonas aromatica* did not occur when O_2_ was present (i.e., oxygen intolerant). Amino acid sequences and predicted structures of NosZ were highly similar among these strains, whereas oxygen-tolerant N_2_O reducers had higher oxygen consumption rates. The results suggest that the mechanism of O_2_ tolerance is not directly related to NosZ structure but rather related to the scavenging of O_2_ in the cells and/or accessory proteins encoded by the *nos* cluster.

**IMPORTANCE:** Some bacteria can reduce N_2_O in the presence of O_2_, whereas others cannot. It is unclear whether this trait of aerobic N_2_O reduction is related to the phylogeny and structure of N_2_O reductase. The understanding of aerobic N_2_O reduction is critical in guiding emission control, due to the common concurrence of N_2_O and O_2_ in natural and engineered systems. This study provided the N_2_O reduction kinetics of various bacteria under aerobic and anaerobic conditions and classified the bacteria into oxygen-tolerant, -sensitive, and -intolerant N_2_O reducers. Oxygen-tolerant N_2_O reducers rapidly consumed O_2_, which could help maintain the low O_2_ concentration in the cells and keep their N_2_O reductase active. These findings are important and useful when selecting N_2_O reducers for bioremediation applications.

## INTRODUCTION

Nitrous oxide (N_2_O) is a potent greenhouse gas and a stratospheric ozone layer destructor [1]. The use of microbial N_2_O reduction has a potential to mitigate N_2_O emissions [2, 3]. This reaction is catalyzed by nitrous oxide reductase (N_2_OR) encoded by the *nos* cluster [4]. N_2_OR is the only known enzyme so far capable of biologically reducing N_2_O to N_2_ and is carried by both denitrifying and non-denitrifying microorganisms [5].

N_2_OR is generally believed to be sensitive to oxygen (O_2_), which may limit the bioremediation application of N_2_OR in a standard aerobic environment. Exposure to oxygen may change the configuration of the copper-based catalytic sites and inactivate N_2_OR [6]. Such inactivation could potentially protect the enzyme from irreversible damage and the production of reactive oxygen radicals upon transient exposure to oxygen [7]. This could also contribute to the sensitivity of N_2_OR to oxygen at the enzyme level. In addition to the effect on the enzyme itself, O_2_ can also influence the transcription of the *nos* cluster. The O_2_-sensing transcription regulators such as FNR and NNR as well as small RNA can suppress the transcription of *nos* [8, 9].

While the impact of O_2_ on microbial N_2_O reduction has been well documented, some denitrifying bacterial strains have been reported to reduce N_2_O in the presence of O_2_ (i.e., aerobic N_2_O reduction) [10]. However, the ecophysiology of aerobic N_2_O reduction remains largely unclear. Questions that remained unanswered include whether the O_2_ sensitivity of N_2_OR is related to their structure and how widely aerobic N_2_O reducers occur in the N_2_OR phylogeny.

There are two distinct clades (Clade I and II) for *nosZ*, the key functional gene of N_2_OR [11]. Genomic differences between the two clades are associated with *nos* cluster organization, translocation pathway, and co-occurrence with other denitrifying genes [12]. Several studies have reported the physiological differences between the two clades. Yoon et al. [13] report that Clade II bacteria (*Dechloromonas aromatica* and *Anaeromyxobacter dehalogenans*) showed high affinities to N2O but lower maximum reduction rates as compared to Clade I bacteria *(Stutzerimonas stutzeri*, formerly known as *Pseudomonas stutzeri* [14], and *Shewanella loihica*). In contrast, Suenaga et al. [3] found that the N2O reduction biokinetics could not be used to distinguish the Clade I bacteria (*St. stutzeri* and *Paracoccus denitrificans*) and Clade II bacteria studied (*Azospira* spp.). Nevertheless, it is still unclear how Clade I and II N_2_O reducers behave in the presence of O_2_.

Therefore, the objectives of this study were to (1) characterize the oxygen sensitivity of various N_2_O reducing bacteria, (2) classify N_2_OR based on their oxygen sensitivity, and (3) examine the relationships between N_2_OR oxygen sensitivity, *nosZ* phylogeny (Clade I *vs*. Clade II), and the predicted N_2_OR structures.

## RESULTS

### Michaelis-Menten kinetics of aerobic and anaerobic N_2_O reduction

By fitting the N_2_O reduction results to the Michaelis-Menten model, we obtained the maximum rate (*V*_max_) and Michaelis constant (*K*_m_) values for various N_2_O reducing strains under aerobic and anaerobic conditions. A wide range of *V*_max_ for nitrous oxide reduction rates was observed. Under anaerobic conditions, bacteria with Clade I N_2_OR generally exhibited faster N_2_O reduction than those with Clade II N_2_OR (**Fig. 1** and **2**). Under anaerobic conditions, *St. stutzeri* TR2 (Clade I N_2_OR) had the highest *V*_max_ (8.37 ± 0.81 μM/s/OD), whereas *G. aurantiaca* T-27 (Clade II N_2_OR) had the lowest *V*_max_ (0.13 ± 0.02 μM/s/OD).

**Figure 1.**
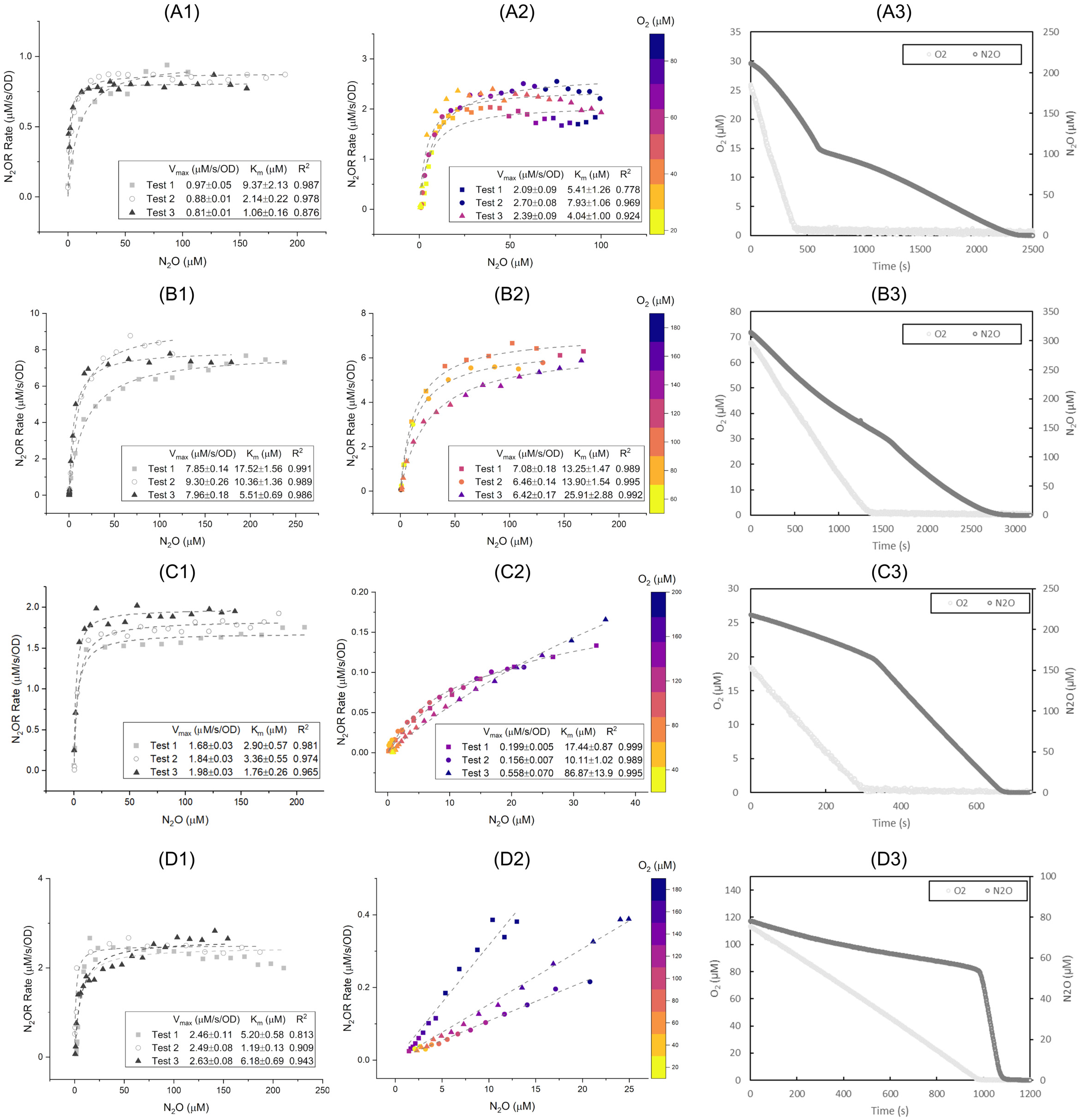
Michaelis-Menten kinetics of (1) anaerobic and (2) aerobic N_2_O reduction, and (3) transition of aerobic into anaerobic N_2_O reduction from (A) *Stutzerimonas stutzeri* ZoBell (B) *Stutzerimonas stutzeri* TR2, (C) *Paracoccus denitrificans* JCM 21484, and (D) *Azospirillum brasilense* Sp7. Curve fitting results were plotted in dashed lines.

**Figure 2.**
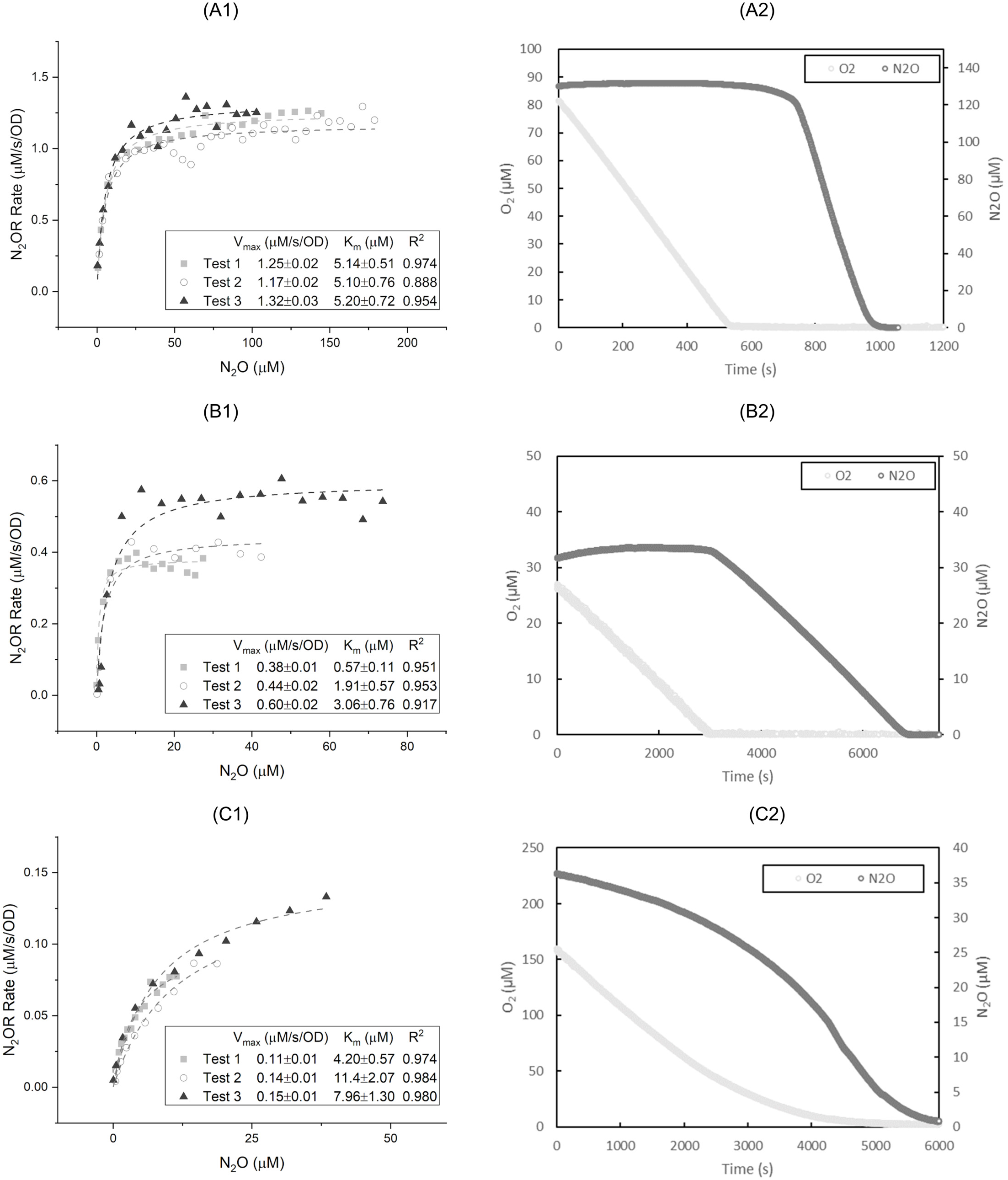
(1) Michaelis-Menten kinetics of anaerobic N_2_O reduction, and (2) the transition of aerobic respiration into anaerobic N_2_O reduction from (A) *Pseudomonas aeruginosa* PAO1, (B) *Dechloromonas aromatica* RCB, and (C) *Gemmatimonas aurantiaca* T-27. Curve fitting results were plotted in dashed lines.

The ability to reduce N_2_O in the presence of O_2_ varied by strain, and there was no overall trend between Clade I and II N_2_OR. For example, *St. stutzeri* TR2 reduced N_2_O under aerobic conditions with the *V*_max_ of 6.65 ± 0.37 μM/s/OD, whereas *Ps. aeruginosa* PAO1 (Clade I N_2_OR) could not reduce N_2_O in the presence of O_2_ (**Figure 2**). Similarly, *D. aromatica* RCB and *G. aurantiaca* T-27 (Clade II N_2_OR) could not reduce N_2_O in the presence of O_2_. *Azospirillum brasilense* Sp7 (Clade I N_2_OR) reduced N_2_O in the presence of O_2_ up to 180 μM; however, its *V*_max_ could not be fitted to the Michaelis-Menten model.

The transition points from aerobic to anaerobic N_2_O reductions (i.e., the change of the slopes between two linear rates) were clearly observed after oxygen was depleted for all tested strains, except for *G. aurantiaca* T-27. For *G. aurantiaca* T-27, the N_2_O reduction rate gradually changed depending on the oxygen concentration (**Figure 2-C2**). In order to further investigate the different oxygen inhibition kinetics observed for *G. aurantiaca*, nonlinear least square fitting with multiple variables was used to determine the inhibition constant (*K*_i_). The non-competitive inhibition model was found to best describe the changing *V*_max_ against various O_2_ and N_2_O concentrations (Figure S1), with a *K*_i_ value of 7.86 ± 1.69 μM O_2_.

The fitted *K*_m_ values for anaerobic N_2_O reduction ranged from 1.85±1.25 μM (for *D. aromatica*) to 11.14±6.04 μM (for *St. stutzeri* TR2). The *K*_m_ values of aerobic N_2_O reduction for *Pa. denitrificans* and *St. stutzeri* TR2 and ZoBell strains did not significantly differ from those of anaerobic N_2_O reduction (Student t-test, *p* >0.05). This indicates that the affinity of Clade I N_2_OR tested did not change with and without the presence of O_2_.

### Classification of oxygen sensitivity of N_2_O reduction

Based on the microsensor analysis, a broad range of N_2_O reduction kinetics was observed under aerobic and anaerobic conditions. As we plot the extrapolated anaerobic and aerobic *V*_max_ values (**Figure 3a**), three distinct types of responses to oxygen were found in the studied strains: 1) Strains with *V*_max_ not affected by oxygen, including *St. stutzeri* ZoBell and TR2, are classified as oxygen tolerant; 2) Strains with much lower aerobic *V*_max_ than anaerobic *V*_max_, including *Pa. denitrificans, A. brasilens*, and *G. aurantiaca*, are classified as oxygen sensitive; 3) Strains that have no N_2_O reduction activity when oxygen is present, including *Ps. aeruginosa* and *D. aromatica*, are classified as oxygen intolerant. NosZ phylogeny seems to be not associated with the classification of oxygen sensitivity. Moreover, the half-saturation coefficients for N_2_O under anaerobic and aerobic conditions agree with previously reported observations. Bacteria harboring Clade II NosZ generally have lower *K*_m_ values than those with Clade I NosZ, suggesting differentiating ecological niches for these two groups of N_2_O-reducing bacteria [13].

**Figure 3.**
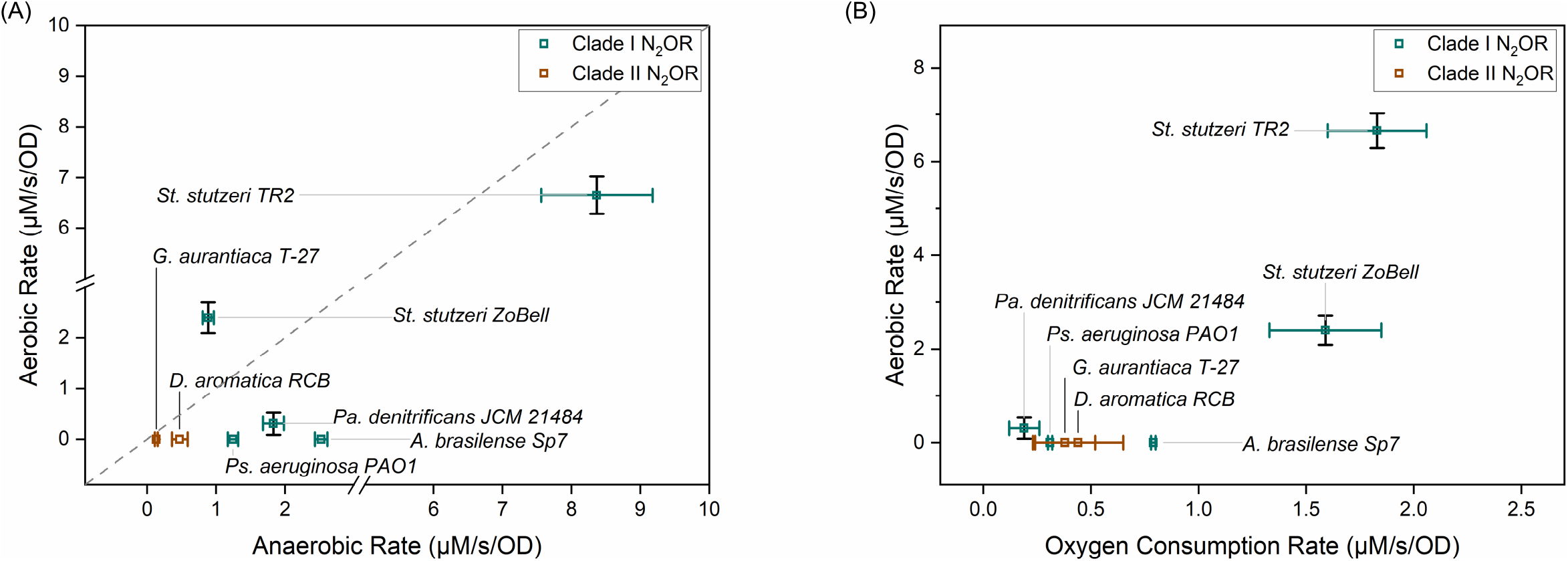
V_max_ for Aerobic N_2_O reduction rates versus (A) V_max_ for anaerobic N_2_O reduction rates and (B) V_max_ for O_2_ consumption of each studied strain.

### NosZ amino acid sequence similarities among the strains

The NosZ amino acid sequences of the strains studied were compared to examine whether the observed differences in oxygen sensitivity originate from the differences in the enzyme structures. Strains investigated in this study cover a variety of classes, including *Alphaproteobacteria* (*A. brasilense* and *Pa. denitrificans*) and *Gammaproteobacteria* (*St. stutzeri* and *Ps. aeruginosa*) for those having Clade I NosZ and *Betaproteobacteria* (*D. aromatica*) and *Gemmatimonadetes* (*G. aurantiaca*) for those having Clade II NosZ. Based on the NosZ phylogenetic analysis Clade I and Clade II NosZ were clearly separated (**Fig. 4**) similar to the previous report [15]. The two *St. stutzeri* strains, both of which showed oxygen tolerant N_2_O reduction, shared a high similarity in the NosZ amino acid sequences (92.6%) (Figure S4). However, *Ps. aeruginosa* PAO1, which showed oxygen-intolerant N_2_O reduction, also had a highly similar NosZ to *St. stutzeri* (77.5% with the ZoBell strain and 79.7% with the TR2 strain). NosZ of oxygen-sensitive N_2_O reducers (*Pa. denitrificans, A. brasilense*, and *G. aurantiaca)* and oxygen-intolerant N_2_O reducers (*Ps. aeruginosa* and *D. aromatica)* were not clustered with each other. In addition, we could not identify amino acid residues that appeared specific to each of the oxygen-tolerant, - sensitive, and -intolerant groups.

**Figure 4.**
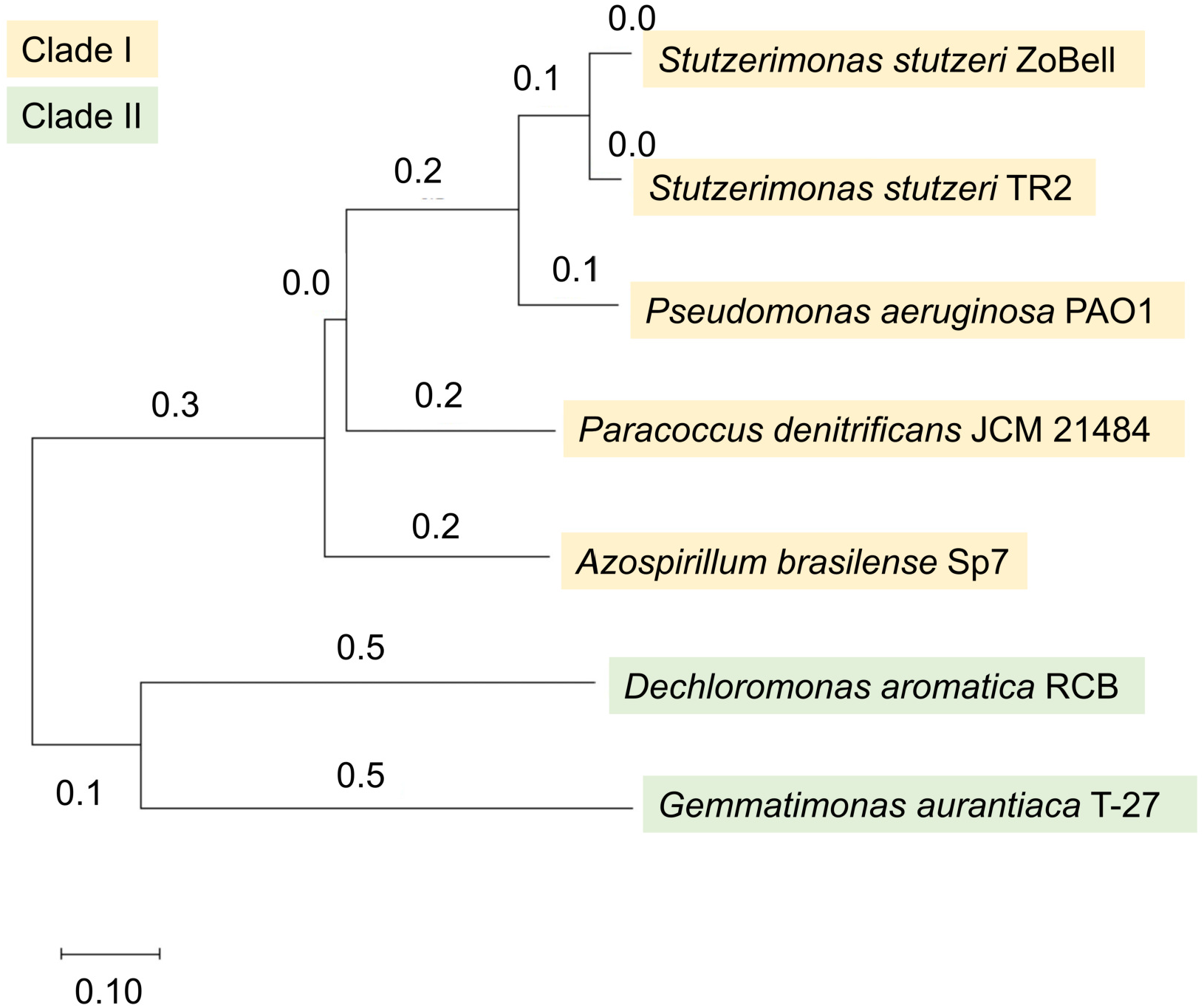
Phylogenetic tree for selected NosZ sequence constructed by Neighbor-Joining method.

Multiple sequence alignment showed that the candidate ligands of Cu_A_ and Cu_Z_ centers were found in all NosZ sequences (Figure S3). The Cu_Z_ catalytic site contains seven histidine ligands which were all conserved in the proposed Cu_Z_ center among Clade I and Clade II (Figure S3). The candidate ligands of Cu_A_ (two cysteines at positions 618 and 622, two histidines at positions 583 and 626, and a methionine at position 629 in *P. stutzeri* NosZ) were also identified in all NosZ (Figure S3).

### NosZ structural similarities

To identify the structural differences between oxygen-tolerant, -sensitive, and -intolerant NosZ, we predicted the enzyme structures based on the NosZ sequences by using Alphafold2 [16] with the ZoBell NosZ [17] as a query structure. We obtained high-confidence NosZ structures, as evaluated based on the sequence coverage and predicted per-residue confidence measure (pLDDT) scores from AlphaFold, with conserved Cu_A_ and Cu_Z_ catalytic domains (Figure S2). Slight structural differences were seen between Clade I and II NosZ as measured by the Dali Z-scores, whereas no differences were seen between NosZ structures from oxygen-tolerant, -sensitive, and -intolerant strains. The Z scores for all Clade I NosZ against the reference ZoBell NosZ were ≥59.8. In addition, the predicted structures for all Clade I NosZ showed the root mean square deviation (rmsd) value of < 2.0 and had no structurally dissimilar amino acid residues of longer than 80 aa to the reference NosZ. In contrast, the Z score for the NosZ of *D. aromatica* and *G. aurantiaca* (Clade II) were 50.8 and 49.3, respectively. Poor matches with the query sequence were obtained for the Clade II NosZ with rmsd values of >2.0 and structurally dissimilar amino acid residues of >80 aa. Most of the structural heterogeneity was observed in the C-and N-terminals.

## DISCUSSION

Biological N_2_O reduction is generally believed to occur under strictly anaerobic conditions. The oxygen sensitivity of N_2_O reduction can be explained by (1) the transcriptional regulation of *nos* and (2) the inactivation of N_2_OR by molecular oxygen. The transcription of *nosZ* can be regulated directly or indirectly by O_2_-sensing transcriptional regulators. For instance, the transcription of *nosZ* is directly regulated by fumarate and nitrate reductase protein (FnrP) in response to oxygen depletion in *Pa. denitrificans* [18]. *Ps. aeruginosa* also has similar FNR-type sensing regulators: The cascading regulation of ANR (anaerobic regulator of arginine deiminase and nitrate reductase) and DNR (dissimilatory nitrate respiration regulator) indirectly controls the synthesis of N_2_OR [19]. Another potential explanation of oxygen sensitivity points to the inactivation of N_2_OR upon exposure to oxygen. The N_2_OR isolated under aerobic and anaerobic conditions exhibited various redox and spin states of copper in active sites. Under limited exposure to oxygen, the enzyme shifted in electron paramagnetic resonance spectra but retained its N_2_O reducing activity. [20] In contrast, aerobic incubation caused loss of copper content and inactivation of the catalytic site. Inactivation of N_2_OR by oxygen was also reported due to irreversible confirmation changes. A sulfur atom binding to the active site of N_2_OR in *St. stutzeri* is lost after exposure to oxygen, leading to the inactivation of the enzyme *in vivo*. [6]

However, these two mechanisms do not explain the occurrence of O_2_- tolerant N_2_O reducers. The cells used for the microsensor experiments were incubated under anaerobic conditions with the addition of nitrite or N_2_O to induce the expression of N_2_OR. One exception is for *G. aurantiaca* T-27^T^. This strain was incubated under aerobic conditions, as *G. aurantiaca* T-27^T^ is an obligate aerobic bacterium that can express *nosZ* in the presence of O_2_ [21]. The same cell cultures were used for aerobic and anaerobic N_2_O reduction rate measurements; therefore, the initial level of N_2_OR expressed in the cells should be the same between the two conditions (i.e., aerobic *vs*. anaerobic N_2_O reduction). Consequently, the transcriptional regulation of *nos* is not contributing to the O_2_ tolerance during N_2_O reduction of each tested strain.

In addition, the structures of NosZ, including the active sites, were highly similar between O_2_-tolerant, -sensitive, and -intolerant N_2_O reducers. Based on the structural similarity and the presence of conserved residues in the active sites, all of the active sites of NosZ and copper cofactors examined most likely receive similar inhibitory effects upon exposure to oxygen [6, 20]. Despite similar N_2_O respiration and bioenergetics in clade I and clade II NosZ, other accessory proteins encoded by the *nos* cluster are expected to function differently [22].

These auxiliary processes could be involved in the maintenance and repair of NosZ, with detailed mechanisms remaining unclear.

Another mechanism that may explain the observed occurrence of O_2_- tolerant N_2_O reduction is the scavenging of O_2_ in the cells. A whole cell assay (as opposed to the assay done with isolated enzymes) was used in this study to calculate the N_2_O and O_2_ consumption rates. When both N_2_O and O_2_ are present, facultatively anaerobic bacteria (e.g., denitrifiers) usually prioritize the respiration of O_2_ over N_2_O because aerobic respiration is more favorable from both bioenergetic and kinetic perspectives [23]. A rapid O_2_ consumption rate can potentially lower the *in situ* O_2_ concentration in the periplasm, where N_2_OR is located. From a simplified estimation shown in the Supplementary Materials, an O_2_ consumption rate of 1 μM/s/OD can cause a significant decrease in O_2_ concentration across cell membranes. When the O_2_ respiration rate is comparable to the O_2_ diffusion rate that replenishes dissolved oxygen in the periplasm, the local oxygen minimum could protect N_2_OR from inhibition in O_2_-tolerant N_2_O reducers. From the tested strains, we indeed observed that bacteria with higher oxygen consumption rates generally have greater oxygen tolerances (**Figure 3b**). A threshold of O_2_ consumption rate could potentially exist, where a lower rate could not emulate the diffusion rate of O_2_ sustaining an anaerobic zone for N_2_OR. Such a protection mechanism could be analogous to the respiration of O_2_ in *Azotobacter* protecting O_2_-sensitive nitrogenase [24].

Our results have some implications for N_2_O removal applications. N_2_O-reducing bacteria, including some of the strains examined in this study, have been used for N_2_O mitigation in natural and engineered systems [2]. For instance, bioaugmentation of *St. stutzeri* TR2 to denitrifying activated sludge has been demonstrated to mitigate N_2_O emissions [25, 26]. *Azospirillum brasilense* strains were also used as a microbial inoculant for N_2_O mitigation in soil. [27] Nevertheless, engineering applications of biological N_2_O mitigation face major challenges including the oxygen sensitivity of N_2_O reduction due to the co-existence and fluctuations of dissolved oxygen and N_2_O concentrations commonly observed in natural and engineered systems. Based on the classification of O_2_ tolerance in this study, kinetic parameters can be used as selection criteria for microorganisms in environmental applications. Oxygen tolerant N_2_ORs were identified only in *St. stutzeri* in this study. Since *St. stutzeri* can reduce N_2_O fast and in the presence of O_2_, this organism is promising for N_2_O bioremediation applications. Besides N_2_O reduction rates, microorganisms with low *K*_m_ values, such as *Pa. denitrificans* and *D. aromatica*, could be useful in scavenging low concentrations of dissolved N_2_O. It is important to note, however, that the kinetics and O_2_ sensitivity of N_2_O reducers can be influenced by environmental factors such as the type of organic carbons [28] and temperature [29]. Therefore, when selecting appropriate N_2_O reducers for engineering applications, their N_2_O reduction kinetics and O_2_ sensitivity should be measured under environmentally relevant conditions.

## MATERIALS AND METHODS

### Bacterial strains

*Stutzerimonas stutzeri* strain TR2 was kindly provided by Dr. Otsubo, Dr. Miyauchi, and Dr. Endo. *Stutzerimonas stutzeri* strain ZoBell (= ATCC 14405) and *Dechloromonas aromatica* strain RCB (= ATCC BAA-1848) were obtained from the American Type Culture Collection (ATCC). *Pseudomonas aeruginosa* PAO1 (= JCM 14847), *Pa. denitrificans* JCM 21484^T^, and *A. brasilense* Sp7^T^ (= JCM 1224^T^) were obtained from the Japan Collection of Microorganisms (JCM). *Gemmatimonas aurantiaca* T-27^T^ (= NBRC 100505^T^) was obtained from Biological Resource Center (NBRC, Kisarazu, Japan).

These strains, except for *D. aromatica* RCB and *G. aurantiaca* T-27^T^, were grown on R2A agar plates amended with 10 mM acetate and 5 mM nitrite under aerobic conditions. After 48 hours of incubation at 30°C, single colonies were picked and transferred to 10 mL of R2A broth with 10 mM acetate and 5 mM nitrite. Each liquid culture was incubated in a sealed tube with an N2 atmosphere at 30°C until harvested during the exponential growth phase. *D. aromatica* RCB was grown on Trypticase soy agar (TSA) supplemented with 5% defibrinated sheep blood under anaerobic conditions at 30°C for 10 days. Single colonies were transferred to 10 mL of R2A broth supplemented with 20 mM lactate and incubated under a 1.39% N_2_O atmosphere (in N_2_) at 30°C until harvested. *G. aurantiaca* T-27^T^ was grown on R2A agar under aerobic conditions. Single colonies were transferred to 10 mL of R2A broth and aerobically incubated at 25°C until harvested. The addition of nitrite inhibited the growth of *G. aurantiaca*, which was expected to have an incomplete denitrification pathway [30].

### Microsensor experiments

Bacterial cultures were harvested during the early to mid-exponential growth phase as determined by OD_600_ measurement. Cultures were washed twice with a sterile 10 mM piperazine-N,N′-bis(2-ethanesulfonic acid) (PIPES) buffer (pH = 7.5) and resuspended in a PIPES buffer supplemented with 10 mM of sodium acetate. The cell suspensions were purged with a gas mix of N_2_O (1.39%, v/v) in N_2_ for 10 minutes to achieve targeted levels of dissolved N_2_O concentrations (300 μM). The cell suspensions were then diluted with PIPES buffer to the desired concentration (~ 10^6^ CFU/mL, OD600 ~ 0.1) and transferred to a double chamber containing mini-stirrer bars (Unisense, Aarhus, Denmark) (Fig. S5). The chamber was capped and placed in a sensor rack with built-in stirrers and submerged in a 30°C water bath. An N_2_O and an O_2_ microsensor (Unisense) were inserted into the chamber via small halls to measure dissolved N_2_O and O_2_ concentrations every second for up to 2 hours or until O_2_ depletion. The N_2_O and O_2_ microsensors were two-point calibrated with zero and saturated solutions (300 μM for N_2_O and 236 μM for O_2_) at 30°C. No cross interference was observed between N_2_O and O_2_ on respective microsensors (Table S1). OD600 of the cell suspension was recorded at the end of each microsensor test. At least three independent microsensor measurements were done for each strain.

The measured concentrations of N_2_O and O_2_ were averaged over time intervals of 100 to 1000 seconds depending on the duration of microsensor tests. This step is useful to minimize the noise generated by the microsensors. Linear rates for N_2_O consumption were extrapolated within each time interval. The Michaelis-Menten plots were then constructed using the rates and corresponding N_2_O concentrations. A nonlinear least square method with the Levenberg-Marquardt algorithm [31] was used for curve fitting on Origin 2021 (Version 9.8.0.200) to determine kinetic parameters including the maximum rate (*V*_max_) and the Michaelis constant (*K*_m_). Similarly, *V*_max_ for O_2_ was linearly extrapolated from O_2_ concentrations measured by the microsensor.

### Bioinformatics and comparative protein structure modeling

The NosZ sequences of the selected strains (GenBank accession numbers WP_011287329, EHY76008, BAM68548, NP_252082, QEL93987, WP_156798935, Q51705 for *D. aromatica* RCB, *St. stutzeri* ZoBell, *St. stutzeri* TR2, *Ps. aeruginosa* PAO1, *A. brasilense* Sp7, *G. aurantiaca* T-27, and *Pa. denitrificans* JCM 21484, respectively) were retrieved from the National Center for Biotechnology Information (NCBI, www.ncbi.nlm.nih.gov). Multiple sequence alignment and phylogenetic tree construction were done using the neighbor-joining method without distance correction by using Clustal Omega (www.ebi.ac.uk/Tools/msa/clustalo/). NosZ structures were predicted through the non-docker implementation of AlphaFold2 version 2.1.1 via the Minnesota Supercomputing Institute (MSI). The NosZ sequence of the selected strains was used as the input with the default prediction parameters to run on a Linux environment. The best-predicted protein models were selected for each sequence and loaded into PyMOL (Schrödinger, Inc., New York, NY, USA). All models were colored based on their pLDDT that are stored in the B-factor fields of the pdb files. All predicted structures were compared against each other using DaliLite.v5 (http://ekhidna2.biocenter.helsinki.fi/dali) [32].

## ACKNOWLEDGEMENTS

We thank Wakako Otsubo, Keisuke Miyauchi, and Ginro Endo for providing *Stutzerimonas stutzeri* strain TR2. We also thank Sujin Yeom and Mike Blazanin for their help with the initial experimental setup and Carrie Wilmot for valuable comments.

This work was supported by the Biocatalysis program and the MnDRIVE Initiative of the University of Minnesota.

